# Monitoring transcription of miR-15a and miR-124 in mesenchymal stem cells from adipose tissue origin differentiated into pancreatic beta-cells

**DOI:** 10.1101/2024.01.18.576237

**Authors:** Hadi Rajabi

**Author notes:** Correspondence to: Dr. Hadi Rajabi (Ph.D.) Koç University Research Centre for Translational Medicine (KUTTAM), Koç University School of Medicine, Istanbul, Turkey.

## Abstract

MicroRNAs are small noncoding pieces of nucleic acid with the potential to control mRNA translation. These sequences participate in the regulation of cell dynamic growth and differentiation. In this study, the expression of miR-15a and miR-124 was monitored in adipose-derived tissue stem cells committed to pancreatic β cells *in vitro* over 28 days. In the current experiment, adipose-derived mesenchymal stem cells were incubated in an induction medium to accelerate differentiation toward endocrine pancreatic lineage for 28 days with a three-stage protocol. To confirm the efficient trans-differentiation of stem cells into pancreatic β cells, we performed a Dithizone staining, a zinc chelating agent, and pancreas-specific hormones (insulin and C peptide) examination via electrochemiluminescence. Real-time PCR analysis was done to assess the expression of miR-15a and miR-124. Dithizone staining confirmed a successful orientation of adipose-derived mesenchymal stem cells into pancreatic β cells indicating with red to strong brown appearance compared to negative control stem cells, indicating insulin positivity in differentiating cells. These effects were prominent by time and reached maximum level at day 28. Concurrently, the expression of both miR-15a and miR-124 was induced and reached a peak expression level at the end stage of the experiment compared to the stem cells (p<0.05).

**Conclusion:** The dynamic of distinct miRNAs, in particular, miR-15a and miR-124 was induced during pancreatic β cells derivation of adipose-derived mesenchymal stem cells.

## 1. Introduction

Diabetes mellitus (DM) is a hyperglycemic condition originating from cellular insulin resistance and/or defects in the dynamics of pancreatic β cells. According to the statistics released by the American Diabetes Association and the WHO, type 1 DM (T1DM) is classified into two main categories such as autoimmune dysfunction and idiopathic forms (1). In addition to pharmacological and conventional approaches, transplantation of cells from pancreatic islets seems to be an appropriate strategy for the alleviation of T1DM-related complications. However, this modality requires lifelong immunosuppression and coincides with remarkable morbidity and negligible mortality (2).

In recent decades, the use of stem/progenitor cells has been increased to restore pancreatic function and elevation of insulin content in the systemic blood system. In line with this progress, regenerative medicine exploits developmental and molecular biology, and stem cell biology as a critical point to address the fate of stem cells after transplantation (3). Considering the remarkable ability of stem cells to proliferate on large scale and trans-differentiate into various cell lineages, these cells are at the center of attention to generate β-cells in diabetic subjects. Thus, β-cells differentiation of stem cells would provide promising unlimited biological sources for reducing diabetes-related complications (4).

Based on a great body of documents, various stem cell sources have been examined to give rise to insulin-producing cells (IPCs) from stem cells (5), induced pluripotent stem cells (iPSCs) (6), and mesenchymal stem cells (MSCs) (7). Considering the lack of ethical issues related to the application of MSCs, these cells are the proper choice to restore pancreatic function compared to embryonic stem cells (ESCs) (8). MSCs can be isolated from different human tissues such as bone marrow (9), adipose tissue (10), and umbilical cord (11). MSCs have the potential to be easily expanded with potent bioactivity to transform into various cell lines (12). Compared to other sources, adipose-derived stem cells (AD-MSCs) have the highest proliferation capacity and even retain multipotentiality after consequent passages (13).

MicroRNAs (miRNAs) are a class of small single-stranded and noncoding RNAs with 17 to 25 nucleotides (14). miRNAs play a fundamental role in the regulation of stem cell differentiation and self-renewal through signaling pathways (15, 16) and the suppression of mRNA translation (17). According to recent studies, the regulatory role of miR-15a and miR-124 was evident in the insulin biosynthesis and β-cell bioactivity (18, 19).

Here, we investigate the possible expression of miRNAs (miR-15a and miR-124) during MSCs differentiation into IPCs. To our knowledge, no enough data is indicating the expression of miRNAs during the differentiation of MSCs-to-IPCs. In this study, we found a change in the transcription of miR-124 and miR-15a during the differentiation of AD-MSCs to IPCs.

## 2. Methods and Materials

### 2.1. Cell culture and expansion

The AD-MSCs cell line was obtained from Royan Institute (Tehran, Iran). Cells were expanded in low-glucose content Dulbecco ‘s modified Eagle ‘s medium (DMEM/LG; Gibco) enriched with 15% fetal bovine serum (FBS, Invitrogen, Carlsbad, USA) in humidified atmosphere at 37°C incubator with 5% CO_2_. Cells at 70-80% confluence were trypsinized using 0.25% Trypsin-EDTA (Gibco) and enzyme activity neutralized by FBS. AD-MSCs at passage 3-6 were subjected to the subsequent analyses.

### 2.2. Cell differentiation into insulin-producing cells

For this propose, the AD-MSCs were pre-treated with three different media; (I) cells were cultivated in DMEM/LG medium containing 5% FBS, 0.5 mM β-mercaptoethanol (Cat no: M6250; Sigma-Aldrich, USA) and 10 mM nicotinamide (Cat no : N0636 Sigma-Aldrich, USA) for 2 days; (II) cells were induced in high-glucose DMEM (DMEM/HG; Gibco, UK) enriched with 2.5 % FBS, 0.5 mM β-mercaptoethanol and 10 mM nicotinamide for next 10 days, and eventually differentiating cells were kept in DMEM/HG medium with 1.5% FBS, 0.5 mM/L β-mercaptoethanol and 10 mM nicotinamide and 10 nM exendin-4 (Cat no: E7144 Sigma-Aldrich, USA) until day 28. The differentiating media were replenished every 2-3 days.

### 2.3. Real time-PCR analysis

In the current experiment, the transcription of miR-124 and miR-15a was monitored during IPC differentiation of AD-MSCs. To this end, an initial number of 3 × 10^5^ AD-MSCs were plated in each well of 6-well plates (SPL) and allowed to reach 70-80% confluency. Thereafter, 3 ml of induction medium were poured to each well and maintained to over a period of 28 days. Following the completion of induction period, total RNA and miRNA contents were isolated from the AD-MSCs on days 0, 7, 14, 21 and 28 by using TRIzol reagent (Cat no: T9424; Sigma-Aldrich, Germany). The quality of extracted RNAs was assessed by using Nanodrop^®^ (Thermo Scientific). cDNA synthesis was conducted with the cDNA synthesis kit (Exiqon, Vedbaek, Denmark) using 50 ng RNA. To designate miRNAs, we used Oligo Primer Analysis Software v. 7. Real-time PCR analysis was carried out using BioMolecular Systems (Model: mic). In analysis was performed in triplicate. In this study, U6 sequence was used as internal control.

### 2.4. Dithizone staining

To confirm the successful differentiation of AD-MSCs to IPCs, we performed Dithizone (DTZ) staining. At the respective time points, DTZ powder (Cat no: D5130; Sigma Aldrich, USA) was liquefied in dimethyl sulfoxide and then sterilized by using 0.2-μm pore size micro-filters. After that, DTZ solution (1% w/v) was added to each cell containing well and incubated for 30 minutes at 37°C. The cells were then precisely washed three times by phosphate buffer saline (PBS). Crimson red stained IPCs were explored under an inverted phase contrast microscope.

### 2.5. Measuring the level on insulin and peptide C by electrochemiluminescence assay

To confirm the functional behavior of IPCs originated from AD-MSCs, we measured the level of insulin and peptide C in the supernatant. After the completion of differentiation protocol, supernatants were collected and the level of insulin and peptide C were measured by Siemens 06602443 Immulite automated bioanalyzer and Roche insulin and C peptide examination kit according to manufactory ‘s instructions.

### 2.6. Statistical analysis

Data are presented in mean ± SD. Statistical analyses were performed by using One-Way ANOVA. P-values<0.05 were considered to be statistically significant.

## 3. Results

### 3.1. miR-124 and miR-15a were up-regulated in AD-MSCs differentiating into IPCs

We used a three steps protocol in order to direct pancreatic differentiation from the AD-MSCs cell line. Based on real-time PCR analysis, we found that the expression of both miRNAs miR-124 and miR-15a was increased in AD-MSCs differentiating into IPCs over a period of 28 days compared to the control group (**Figure 1**). We found that first 7-day incubation with differentiation medium did not change the transcription level of miR-124 and miR-15a while these values were increased two-week after AD-MSCs induction toward IPCs. In this regard, the level of miR-124, but not miR-15a, reached significant levels (*p*<0.05). The level of miR-124 was reached maximum level 28 days after treatment (p<0.05) (**Figure 1**). Similar to miR-124, miRNA-15a increased significantly at the endpoint of experiment. These data demonstrated an upward trend in the level of miR-124 and miR-15a during differentiation of AD-MSCs toward IPCs.

**Fig. 1.**
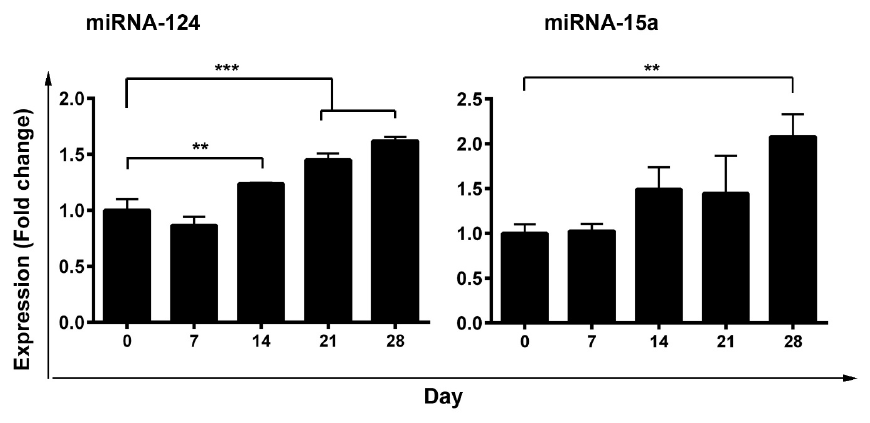
Monitoring the expression of miR-15a and miR-124 during the differentiation of AD-MSCs into IPCs detected by real-time PCR. Data are presented as mean ± SD. All experiments were conducted in triplicate. One-Way ANOVA and post-hoc analysis; ^**^p<0.01 and ^***^ P<0.001.

### 3.2. DTZ staining confirmed AD-MSC-IPC differentiation

Pancreatic islets located at endocrine part contain a large amount of zinc components in comparison with other tissues. As a matter of fact, DTZ is used as a zinc-binding substance to discover the existence of these compounds inside IPCs, showing the level of trans-differentiation (**Figure 2**). Bright field microscopic imaging showed the crimson red stained clusters in differentiating AD-MSCs after 28 days compared to the non-treated control AD-MSCs (**Figure 2**). These data showed that 28-day incubation of AD-MSCs with differentiation medium caused to accumulation of zinc components inside IPCs, showing the successful phenotype acquisition.

**Fig. 2.**
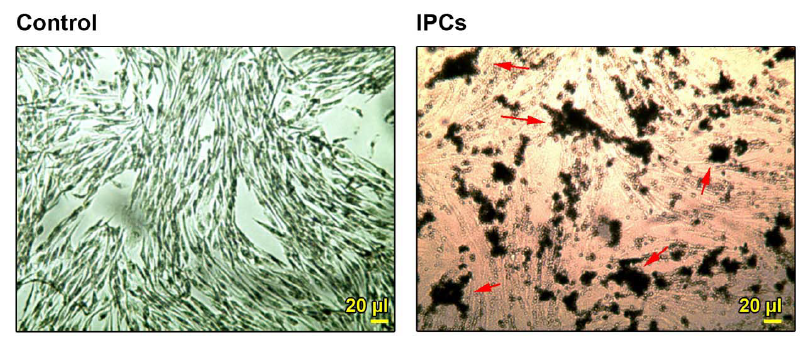
Detection of insulin in differentiating cells toward IPCs by DTZ staining after 28 days. The existence of red to strong brown granules showed the insulin peptide inside the cells. The control stem cells were negative to DTZ staining.

### 3.3. IPCs had potency to secret insulin and C peptide

The ability of IPCs to secret C peptide and insulin could show the functional behavior of target cells after induction of phenotype acquisition. Results from electrochemiluminescence panel showed the increase of insulin (p<0.05) and C peptide (p<0.01) in the supernatant of differentiating cells compared to the control group (**Figure 3**). These results showed the functional behavior of differentiated cells from AD-MSCs sources.

**Fig. 3.**
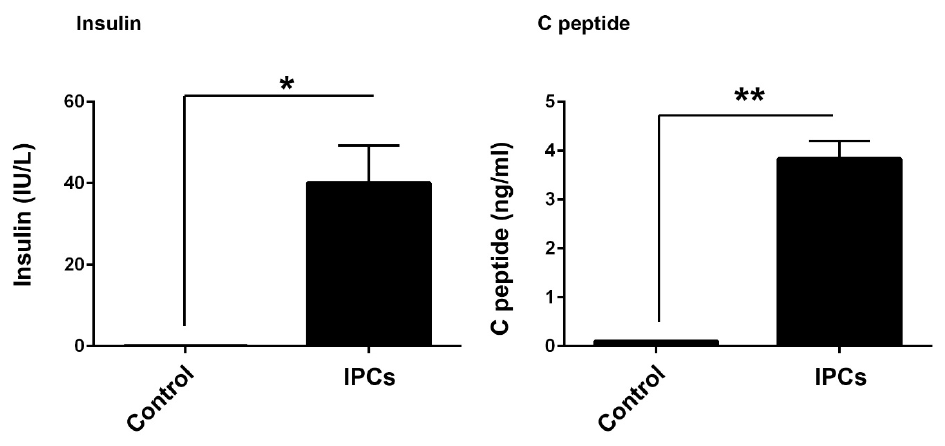
Detection of released C peptide and insulin in AD-MSCs after trans-differentiation into insulin producing cells. Biochemical analysis revealed the significant differences in the levels of C peptide and insulin in culture medium of differentiating cells compared to the control non-treated cells. Student t test. ^*^ P<0.05 and ^**^P<0.01 (n=3).

## 4. Discussion

Stem cells are known as natural cell resource for the regeneration of injured tissues. Compared to the current approaches, stem cell therapy paves a way with promising results to alleviate diabetes-related complications(20). It was found that these cells have potential to produce IPCs after administration to pancreatic tissue (3). According to a great body of documents, various stem cell types were used to produce IPCs such as ESCs, MSCs etc.(21),(13). Even, there was a report on the potency of induced pluripotent stem cells (IPS) in orientation toward IPCs (22). In this regard, MSCs have been introduced as a suitable cell resource for application in different pathologies(23). To acquire novel cell phenotype, different intracellular signaling pathways, effectors and genetic elements are involved in favor of trans-differentiation.

However, there is no consolidated agreement about regulatory mechanism that dictate stem cells trans-differentiation to various cell types especially IPCs(24, 25), (26). Considering the regulatory effect of miRNAs in cell bioactivity, the analysis of miRNAs profile could shed a light for giving optimum stem cells differentiation rate (27). In support of this statement, experiments showed that the inhibition of Dicer 1, an RNAase endonuclease that synthesizes miRNAs, contributes to modification of islet structure, decrease of β-cells and insulin secretion in response to glucose (28). As a matter of fact, selection of appropriate miRNAs could be helpful in detection, dynamics and differentiation capacity of stem cells to IPCs. Previously, Wei and his colleagues explored expression patterns of miR-375, -7, -146a and -34a during the ESCs differentiation into IPCs. Based on their results, a diverse expression pattern was found, showing the distinct effect of each miRNA in the dynamic of stem cells toward IPCs (5).

To our knowledge, there are few reports related to critical effects of miR-124 and miRNA-15a dynamics during AD-MSCs to IPCs orientation. In line with this statement, we aimed to explore the expression of both miRNAs over a period of 28 days. Our data showed that the expression of both miR-124 and miRNA-15a increased during IPCs differentiation of AD-MSCs which coincided with zinc elements and cell potency to deliver insulin and C peptide. Consistent with our results, Sebastiani et al. monitored the expression of eighteen miRNAs especially miR-124a from miRNA family and noted the induction of this miRNA inside IPSCs committed to IPCs. They also revealed that the transcription of other miRNAs such as miR-135a, miR-138, miR-149 and miR-375 was induced while the miR-31, miR-127, miR-143, miR-373 profoundly suppressed (29). Empirical studies found a close relation of miR-15a and miR-124 with insulin biogenesis inside β cells. In a recent study conducted by Sun and co-workers, it was found that miR-15a could inhibit uncoupling protein-2 belongs to mitochondrial inner membrane carriers that prohibit insulin secretion (18). The increase of miR-15a could decrease the inhibitory effect of uncoupling protein-2 on insulin secretion. On the other hand, the induction of miR-124 takes part in the development of pancreatic β-islets via engaging fork head box protein A2 (19). In consistent with our data, both miR-124 and miRNA-15a were increased during IPCs generation. The analysis of these miRNAs possibly could be monitored to address the successful differentiation toward IPCs.

We are aware of some limitation regarding the current experiment. We suggest ongoing experiment will be conducted to monitor the expression of multiple miRNAs at the same time in prolonged time period. It is mandatory to find the possible relationship between above-mentioned miRNAs with pancreatic specific factors.

